# Predator Odor Stressor, 2,3,5-trimethyl-3-thiazoline (TMT): Assessment of stress reactive behaviors during an animal model of traumatic stress in rats

**DOI:** 10.1101/2023.11.07.566080

**Authors:** Laura C. Ornelas, Joyce Besheer

**Author notes:** Correspondence: Joyce Besheer, Ph.D., Professor, Bowles Center for Alcohol Studies Department of Psychiatry, University of North Carolina at Chapel Hill Chapel Hill, NC 27599-7171.

## Abstract

Animal models utilizing predator odor stress are important in understanding implications for post-traumatic stress disorder. 2,5-dihydro-2,4,5-trimethylthiazoline (TMT) has been used to measure stress reactive behaviors during TMT exposure, indicative of stress coping behaviors. In addition, long-term consequences of stress including contextual-induced stress memory, anxiety-like and hyperarousal behaviors, and subsequent increases in alcohol self-administration can also be examined after TMT exposure.

In this article, we describe the TMT exposure protocol used in our lab and how we measure different stress-reactive behaviors that rats engage in during the TMT exposure. Rats are placed in Plexiglas chambers that contains white bedding on the bottom of the chamber and a metal basket in the top right corner that contains a filter paper on which 10 µL of TMT is pipetted onto. During the 10 min exposure, rats can move around the chamber freely. Exposures are recorded by a video camera for later analysis.

During TMT exposure, rats engage in a variety of stress-reactive behaviors, including digging and immobility behavior. These are two distinctly different types of stress-induced behavioral coping strategies to measure individual differences in stress responsivity. To examine individual differences, we group rats into TMT-subgroups based on time spent engaging in digging or immobility behavior. We calculate a digging/immobility ratio score in which we divide the total time spent digging by the total time spent immobile. A cut-off strategy is used such that rats with a criterion ratio score below 1.0 are classified as TMT-1 (i.e., low digging/high immobility; greater passive coping) and rats with a ratio score above 1.0 are classified as TMT-2 (i.e., high digging/low immobility; greater active coping).

In the present paper, we provide a detailed description of the TMT exposure protocol and step-by-step process in evaluation of stress-reactive behaviors.

**Basic Protocol 1:** Predator Odor Stressor Exposure using TMT

**Basic Protocol 2:** Description of Stress-Reactive Behaviors during TMT Exposure and Formation of TMT-subgroups

## INTRODUCTION

Animal models have become increasingly important in stress research to examine behaviors that can inform our understanding of clinical PTSD symptoms. Exposure to a predator odor stressor has been shown to be a valid and reliable animal model to examine symptom profiles of PTSD, as well as lasting consequences to traumatic stress (Cohen, Kozlovsky, Alona, Matar, & Joseph, 2012; Staples, 2010; Verbitsky, Dopfel, & Zhang, 2020). For example, experiencing intrusive distressing memories of a traumatic event(s) is a DSM-5 diagnostic criteria for PTSD (APA, 2013). Animal models of predator odor stress, such as bobcat urine produces contextual avoidance to an odor-paired chamber in rats (Albrechet-Souza & Gilpin, 2019; Weera, Schreiber, Avegno, & Gilpin, 2020) and exposure to the synthetically produced predator odor 2,5-dihydro-2,4,5-trimethylthiazoline (TMT; an extract of fox feces), produces contextual stress responses, as evidenced by increased freezing behavior during re-exposure to the exposure context (Ornelas, Tyler, Irukulapati, Paladugu, & Besheer, 2021; Ornelas, Van Voorhies, & Besheer, 2021; Schwendt et al., 2018; Tyler, Bluitt, Engers, Lindsley, & Besheer, 2022; Tyler, Weinberg, Lovelock, Ornelas, & Besheer, 2020).

Predator odor stressor exposure can also be used to investigate the complex interaction between increased alcohol consumption and stress, which may help in understanding the neurobiology associated with comorbid PTSD and alcohol use disorder (Ornelas, Tyler, et al., 2021; Weera et al., 2020). For example, several studies have shown increased alcohol drinking (Alavi et al., 2022; Albrechet-Souza, Schratz, & Gilpin, 2020; Makhijani, Franklin, Van Voorhies, Fortino, & Besheer, 2021) or alcohol relapse-like behavior (Becker, Lopez, King, & Griffin, 2023; King & Becker, 2019) following exposure to varying predator odor stressors, including TMT. Importantly, not all individuals who experience a traumatic event develop PTSD. There may be individual differences in behavioral responses to stress, which may contribute, in part, to susceptible and resilient populations (Center for Substance Abuse, 2014).

Using the TMT exposure procedure outlined in this current protocol, we are able to focus on individual differences in stress reactivity behavior (i.e., coping behaviors), by classifying rats into specific phenotype groups. Using these behavioral phenotype groups, we can then assess how engagement in different coping strategies during stress exposure may lead to or mitigate against lasting consequences of stress (for example, contextual stress memory, anxiety-like behavior, hyperarousal, or alcohol drinking). For example, we have found that rats, specifically female rats, that engage in greater digging behavior versus immobility behavior during TMT exposure show increased alcohol self-administration after stress (Ornelas, Tyler, et al., 2021). Therefore, by quantifying behaviors that occur during the stressor, we can then assess the relation between stress reactivity phenotypes and the stressor-induced neurobiological and behavioral changes.

The current exposure protocol takes advantage of observing and quantifying the behaviors that the rats engage in *during* the TMT stressor exposure. Therefore, one can quantify a multitude of stress-reactive behaviors including avoidance, darting, grooming, midline crossings, defensive digging and immobility behavior. However, we specifically focus on defensive digging and immobility because they capture two distinctly different types of coping behaviors (digging = active, immobility = passive) that we have found useful to characterize individual differences in stress responsivity.

In this article we will provide detailed description of the specific TMT exposure protocol we utilize to examine individual differences in stress-reactivity in rats.

## STRATEGIC PLANNING

Below are necessary procedural, planning and experimental design details that are important to follow prior to starting a TMT exposure.

### Animals

1. Upon arrival to the vivarium, rats (Long-Evans) are double housed by sex with ad libitum food and water. Our published reports have used adult Long Evans rats and this protocol is based on Long Evans rats; however, this does not preclude the use of other rat strains.
2. Housing procedures after arrival to the vivarium are dependent on experiments. For example, if there are surgical manipulations or certain behavioral procedures, rats may need to be single housed (Ornelas, Van Voorhies, et al., 2021) (Ornelas, Tyler, et al., 2021). However, it is also an option to double housed rats. Housing conditions will likely need to be factored into experimental planning discussions. Rats are handled for five days prior to the start of any experimental procedures.

### Apparatus

3. Plexiglass chambers used for water and TMT exposures were custom built by the University of North Carolina at Chapel Hill Instrument Shop.
4. The length of the back wall (45.72 x 20.32 cm; KYDEX® sheet material) of the chambers is opaque white with two opaque black side walls (left wall with latch for lid: 19.05 x 16.51 cm; right wall: 20.32 x 16.51 cm; KYDEX® sheet material) and a clear front wall (45.72 x 20.32 cm; Plexiglass) to allow for video recoding. The bottom of the exposure chamber is clear (45.72 x 20.32 cm; Plexiglass). To secure the exposure chamber and prevent rats from escaping, a removable clear Plexiglass lid (50.8 x 17.78 cm; Plexiglass) is inserted into the top of the chamber. *(See Figure 1 for material breakdown; see Figure 2 for full schematic representation)*.
5. A metal basket (2.54 x 2.54 cm; ∼17.78 cm above the floor; Plumb Pak by Keeney Sink Basket Strainer) is hung on the front right corner of the front side wall corner (see Figure 1 for representative image) from a metal holder (7.62 x ∼6.35 cm custom build by UNC-CH Instrument Shop). This basket holds a piece of filter paper (cut from GE Healthcare LifeSciences Whatman^TM^, Filter Paper; triangle shape, 3.81 x 3.81 x 3.81 cm) on to which water or TMT (10µL) will be is pippetted later (Step 3 under Water/TMT Exposure steps). The filter paper is inaccessible to the rat. *(See Figure 1)*.
6. At the bottom of the exposure chamber, approximately 600 mL or 130 g of white bedding (Shepherd, Alpha-dri®; virgin paper pulp cellulose that is completely clean and virtually free of contaminants) is added to the bottom of the exposure chamber prior to the animal being placed in the chamber. Bedding is spread eveningly throughout the bottom of the chamber.
7. To prevent TMT from contaminating water exposure chambers, we have separate chambers that are used for water and TMT exposure. Water and TMT chambers are identical in physical dimensions.
8. Appropriate cleaning procedures are essential when using predator odor compounds to prevent contamination or lingering predator odor scent. After the completion of all TMT exposures, water exposure chambers are cleaned first and TMT chambers are cleaned second. However, the step-by-step cleaning procedure is the same for water and TMT exposure. First, bedding is discarded and all lids are removed. Metal baskets are removed from the chamber and filter paper is discarded. Metal baskets and holders are soaked in 3% bleach solution for at least 24 hr then air dryed. For good measure, metal baskets and holders for water and TMT should be soaked seperatedly in bleach solution. Chamber lids are sprayed with Peroxigard^TM^ One-Step Disinfectant Cleaner and Deodorizer (Virox® Technologies Inc.) with a 1-minute contact time. Chamber are first sprayed with water then extensively sprayed with Peroxigard^TM^. After at least 1 min of contact time, chambers are rinsed with water then air dryed on drying rack. Trash that contains TMT-exposed material (bedding, filter paper, pipette tips, aliquot containers) is removed from the procedure room and the animal facility.

### Shelving Unit

1. During water and TMT exposure, the chambers are placed on a shelving unit during video recording. The shelving unit is constructed from UNC-CH Instrument Shop. It is made out of KYDEX® sheet material.
2. Dimensions of the shelving unit are 76.2 x 111.76 cm with six comparments for chambers (see *Figure 2A* for schematic representation of shelving unit).
3. Shelving units are placed on a counter top in the procedure room in front of a camera and tripod for recordings (∼ 4ft (121.92 cm)).
4. This shelving unit allows for recording of one to six rats at a time during one single exposure.

### Predator Odor

1. 97% purity 2,5-dihydro-2,4,5-trimethylthiazoline (TMT) is purchased from SRQ Bio (Sarasota, FL) in 1 gram glass vials. TMT is a clear light yellow, synthetic compound derived from from fox feces.
1. Aliquots of TMT (150 µL) are made upon arrival and stored at -20°C until ready to be used. When pipetting TMT, always do so under a fume hood. In addition, any PPE material (i.e., gloves) that comes into contact with TMT should be discarded immediately to prevent contamination of TMT on surfaces. Furthermore, at the end of all TMT exposures, all TMT-contaminated materials must be removed from the procedure room and animal facilities.

### Environment

1. Predator odor exposures should occur in procedure rooms that allow for ventilated and circulated air flow. In addition, rooms should be equipped with a fume hood when pipetting TMT odor.
2. All protocol steps for predator odor exposure including exposure and cleaning should occur in the procedure room to prevent contamination of odor throughout facilities.
3. Importantly, individuals should be cautious about using the procedure room after a TMT exposure as the room needs time for the predator odor to be ventilated out of the room, as lingering TMT could affect the behaviors of other animals that enter the room. This is dependent on flow rate of air in and out of procedure room.

**Figure 1.**
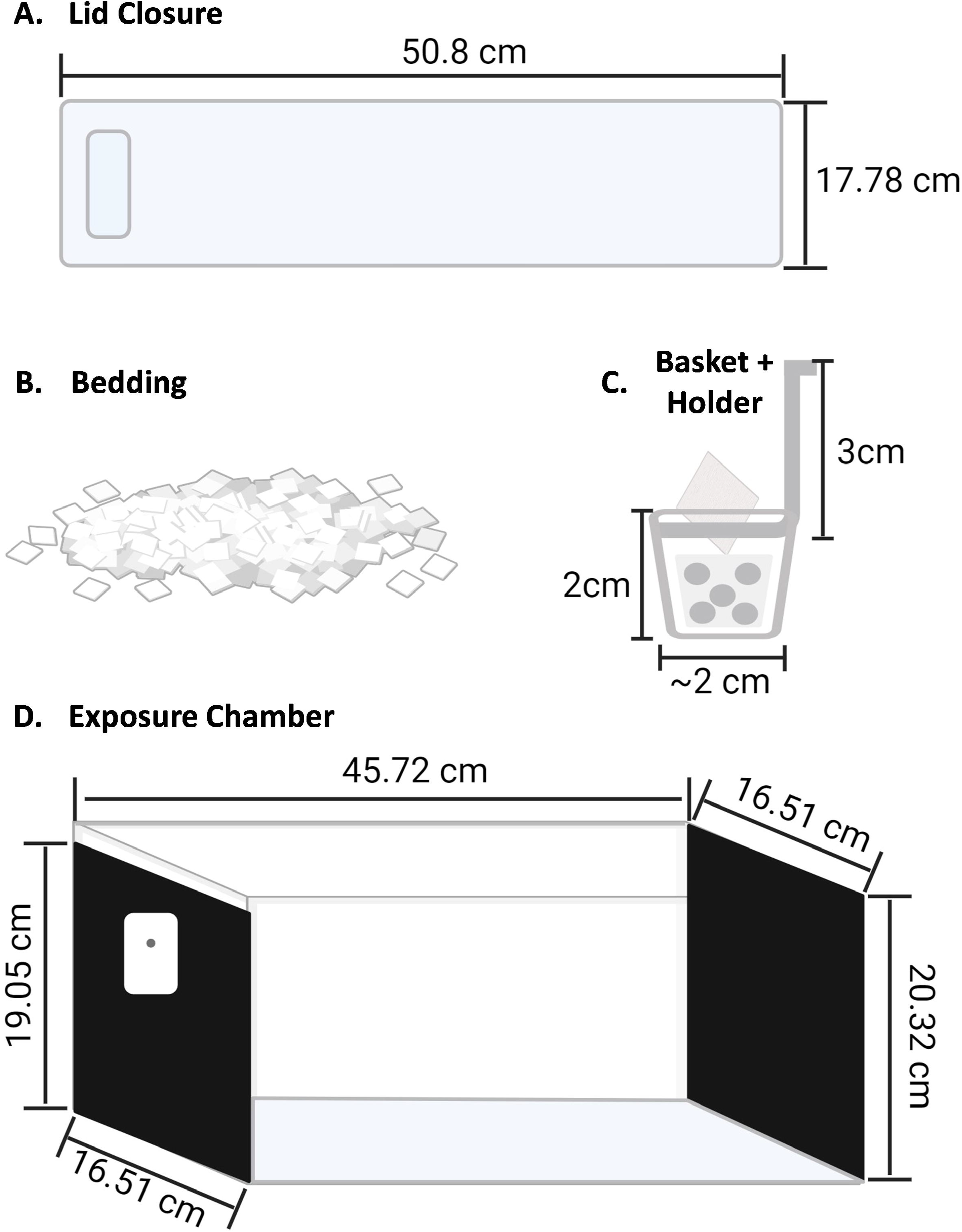
Description of exposure chamber materials. (**A**) Lid closure, (**B**) Alpha-Dri bedding added to bottom of chamber, (**C**) basket holder and basket that holds filter paper for water or TMT and (**D**) outline of exposure chamber. Created with BioRender.com.

**Figure 2.**
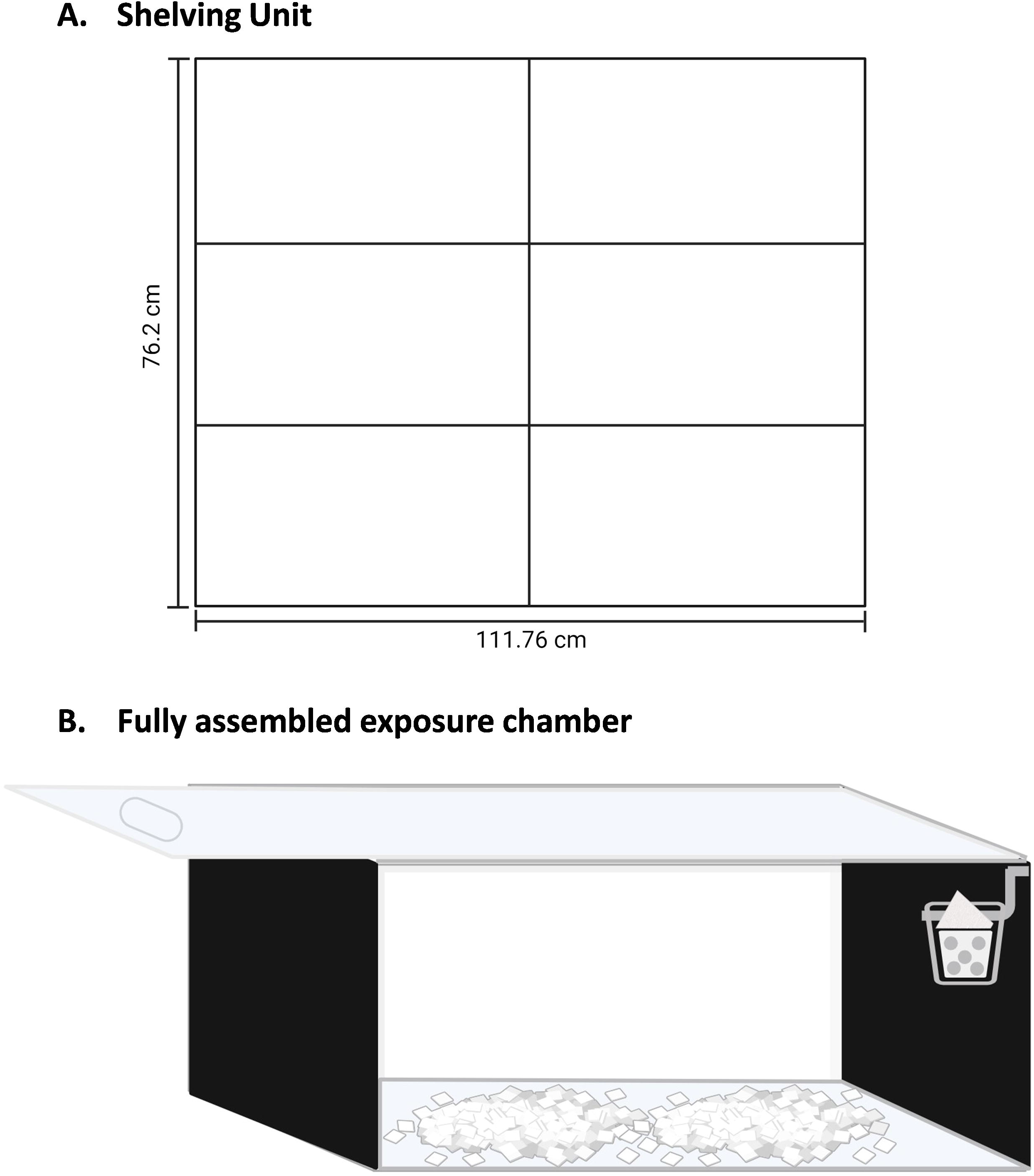
Description of shelving unit and schematic representation of fully assembled exposure chamber. (**A**) Six-slot shelving unit used to hold exposure chambers during recordings. (**B**) Illustration of fully assembled exposure chamber with closing lid, bedding, basket and holder with filter paper. Created with BioRender.com.

## BASIC PROTOCOL 1

**Basic Protocol 1 Title:** Predator Odor Stressor Exposure using TMT

### Introductory paragraph

This protocol aims to explain how we perform a TMT exposure procedure in the laboratory. Prior to exposure, animals are randomly assigned to water or TMT exposure groups. All necessary exposure materials and equipment (detailed description below) are set up in a well-ventilated procedure room prior to transporting rats into the room. After rats are transported into the procedure room, water or TMT (10 µL) is pipetted onto filter paper that is placed in the metal baskets in exposure chambers under a fume hood. Each rat is individually placed in the assigned chamber, then the lid is secured and the chamber is placed on the shelving unit where the exposure will be video recorded. Exposures last for 10 minutes then rats are removed from their chambers and returned to their home cage and returned to the vivarium.

### Materials

Cart/transportation for rat cages from vivarium to procedure room
Rats
Water/TMT Exposure Chambers (see strategic planning section for description, measurements and included materials)
Trimethylthiazoline (TMT; 97% purity, BioSRQ)
µL Pipette with pipette tips
Stop watch/timer
Shelving unit(s) for exposure chambers Video camera
Memory Card for camera
Tripod
Experimental data sheet that includes rat number, treatment group, exposure time, layout key of exposure order on shelving unit, date, name of experimenter)
Pen
Tape
Trash bin
Peroxiguard/Cleaning solution
Paper towels

### Protocol Steps

#### Before transporting animals to procedure room for water/TMT exposure

1. Prepare water and exposure chambers – add bedding (600 mL) to bottom of the chamber, place metal basket on front corner with filter paper. Fasten lid to top of chamber.
2. Place shelving units, which hold exposure chambers for recordings, on procedure tables.
3. Set up Tripods for recordings with video camera in front of shelving unit to record up to six exposure chambers during one exposure session.

*Note: Recording camera needs to be fully charged and there needs to be enough room on memory card for amount of exposure that will be recorded. If the video camera runs out of battery or space on memory card while recording an exposure video, that will result in lost data for behavioral observation*.

4. You will start with water exposure first. Place water exposure chambers under fume hood with lid half way open to allow room for placement of the rat in the chamber. Placement under fume hood allows the same transfer from cage to chamber procedure for control and TMT rats.
5. Tape experimental run sheet with water/TMT group assignments to fume hood or in the procedure room to follow exposure order and key for chamber odor on shelving unit.
6. To preserve camera battery life, turn on camera immediately before the start of the exposure and turn off after the exposure is finished.

#### Water/TMT Exposure

1. Transport rats to procedure room. Only transport rats that will undergo exposure for this specific exposure session. *Note: It is not necessary to habituate rats to transport and the procedure room prior to exposure. However, when transporting rats to procedure room, reducing the amount of noise and disturbances during transport is important to prevent confounding effects during stress exposure*.
2. Open lid for all exposure chambers under fume hood.
3. Pipette water (10 µL) onto filter paper for all chambers.
4. Turn camera on. Press record button.
5. Open rat cage for first rat. Retrieve rat and place in assigned exposure chamber.
6. Close lid to exposure chamber, and carry the chamber to the shelving unit and place the chamber in designated spot on shelving unit. *Note: Depending on the layout of the room, make sure to not accidently hit camera or tripod while recording. When placing rats onto shelving unit, walk around the tripod and camera with caution*.

7. Repeat for each rat until the last rat has been placed in the chamber and on shelving unit (*see Figure 3 for exposure illustration*).
8. Start timer for 10 min.
9. Leave procedure room.
10. At end of exposure, enter procedure room.
11. Stop recording and turn camera off.
12. Remove each chamber individually place rat back into its home cage.
13. Return rats to vivarium.

**Figure 3.**
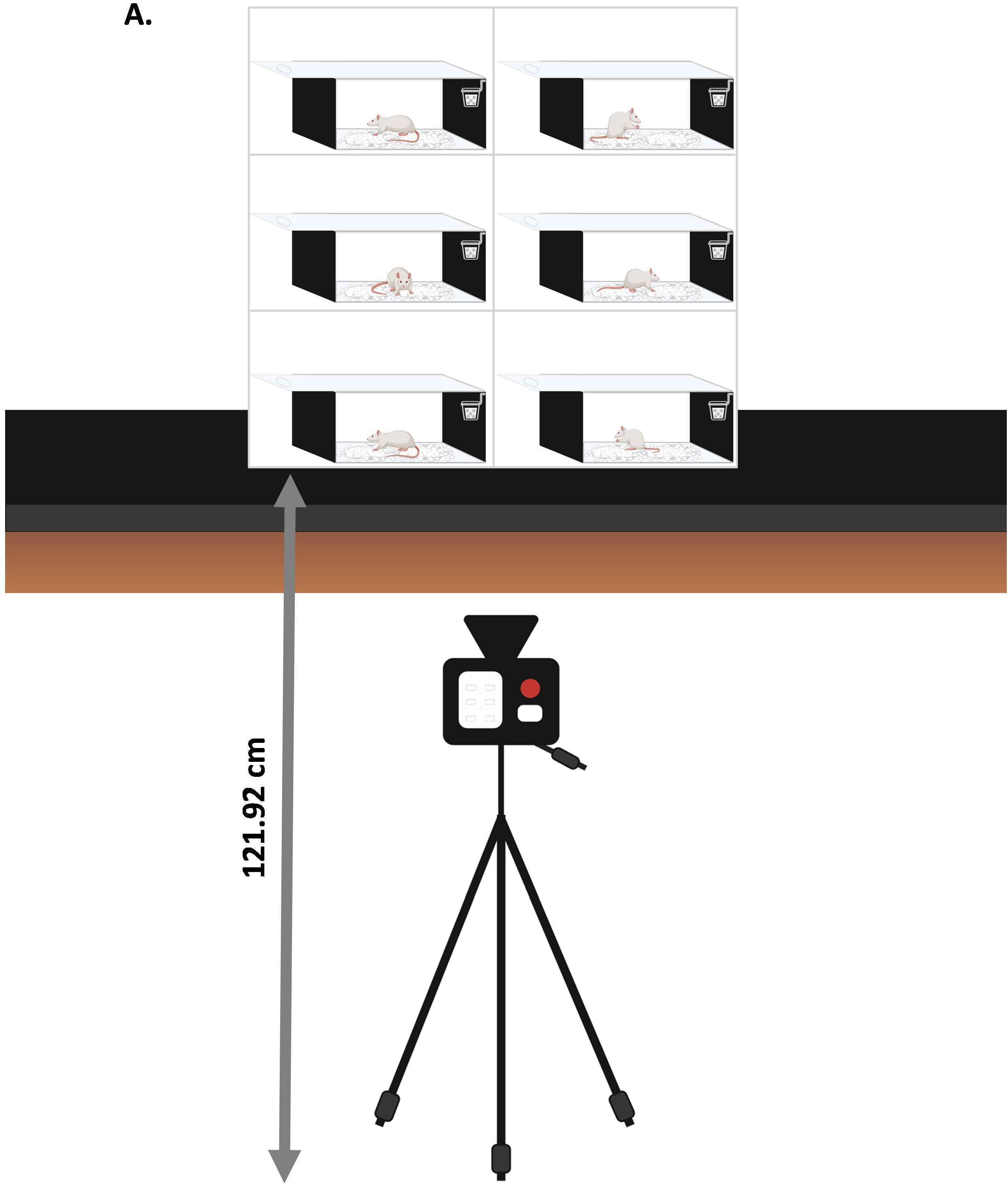
Illustration of exposure set-up. (**A**) Recording camera is approximately 121.92cm (4ft) directly in front of shelving unit and exposure chambers. Up to six exposure chambers can be placed on the shelving unit during recordings. Created with BioRender.com.

## BASIC PROTOCOL 2

**Basic Protocol 2 Title:** Description of Stress-Reactive Behaviors during TMT Exposure and Formation of TMT-subgroups

### Introductory paragraph

During the TMT exposure, we can quantify a multitude of stress-reactive behaviors including avoidance, darting, grooming, midline crossings (indicative of locomotor movement), defensive digging and immobility behavior. For the purpose of this current protocol and description of how our lab evaluates individual differences in stress reactivity, we focus on defensive digging and immobility/freezing behavior. Protocol 2 focuses on 1) a detailed description of defensive digging behavior and immobility/freezing behavior during TMT exposure and 2) formation of TMT-subgroups to examine individual differences in stress reactivity.

### Materials

SD Memory Card for camera

USB 3.0 SuperSpeed SD/Micro SD Card Reader (transfer video files from SD memory card to computer)

Adobe Premier Pro (to convert video files if needed for behavioral analysis software)

Behavioral analysis software (may be helpful but not required as behaviors can be scored manually)

### Protocol Steps

#### Defensive Digging Behavior

1. To allow for digging behavior to occur during TMT exposure, bedding must be present in the exposure chamber (see *Basic Protocol 1*)
2. During the TMT exposure, engagement in digging behavior typically occurs at the start of the exposure; however, the pattern of digging behavior is variable and dependent on each rat.
3. It is important to note, not all rats will engage in digging behavior. The amount of digging behavior will vary between each rat. This is a unique factor about this specific stress-reactive behavior that makes it advantageous to study individual differences.
4. Defining what constitutes as digging behavior during TMT exposure was first developed using De Boer & Koolhass (2003) definition of defensive burying:

a. Rodents will alternate between forward pushing and shoveling movements with their forepaws and head directed at a threat (i.e. TMT) (De Boer & Koolhaas, 2003).
5. During the TMT exposure, the rats will actively thrust bedding material to the corner of the chamber under the metal basket where the TMT is located.
6. *Note: exploratory behavior (i.e. sniffy and inspecting bedding with nose or forepaws) during exposure does not constitute as digging behavior.
7. The TMT basket is inaccessible to the rat; therefore, the rat is unable to “bury” the threatening stimuli (i.e. TMT). As a consequence of the burying, rats will build a large corner pile of bedding material throughout the TMT exposure.
8. Digging behavior is quantified manually and analyzed using ANY-maze ™ Video Tracking System (Stoelting Co. Wood Dale, IL, USA); however, any behavioral tracking software that is preferred can be used to quantify digging behavior.

#### Immobility/Freezing Behavior

1. During the TMT exposure, engagement in immobility behavior is also variable between each rat.
2. However, engagement in this stress-reactive behavior typically occurs towards the middle to end of the TMT exposure.
3. Immobility is operationally defined as lack of movement for more than 2 seconds as assessed using ANY-maze ™ Video Tracking System (Stoelting Co. Wood Dale, IL, USA) (Makhijani et al., 2021; Ornelas, Tyler, et al., 2021; Ornelas, Van Voorhies, et al., 2021; Tyler et al., 2022; Tyler et al., 2020)
4. *Note: immobility likely captures both inactivity and freezing behavior.

#### TMT-Subgroup Classification

1. Subgrouping rats based on stress-reactive behaviors allows for examination of individual differences in stress response.
2. Using the TMT exposure described in *Basic Protocol 1* and behaviors previously described, we first quantify for each rat, the total time a rat engages in digging and immobility behavior.
3. We classify rats into TMT-subgroups using a digging/immobility ratio score that is calculated by dividing the total time spent digging by the total time spent immobile.
4. A cut-off strategy is used such that rats with a criterion ratio score below 1.0 were classified as TMT-1 and rats with a ratio score above 1.0 were classified as TMT-2 (Ornelas, Tyler, et al., 2021; Ornelas, Van Voorhies, et al., 2021).

## COMMENTARY

### Background Information

There are many advantages of using 2,5-dihydro-2,4,5-trimethylthiazoline (TMT) as a predator odor stressor. It activates a hardwired “learned-independent system” shown to induce innate fear and defensive behaviors (Rosen et al., 2008; Rosen, West, & Donley, 2006). Additionally, it is considered a less invasive model compared to other animal models such as stress-enhanced fear learning or conditioned fear models that utilize shock stressor, single-prolonged stress that uses restraint, swim and ether anesthesia, and social defeat stress in which animals are repeatedly exposed to a social defeat situation.

Another advantage of predator odor stressor models in general, is the ability to examine individual differences in behavior *during the stressor*, as an index of stress reactivity, and then classifying animals into specific phenotypic groups. Previous studies have used characterization methods that focus on grouping animals based on behavioral changes that occur*after* stress.

For example, animals have been sub-grouped into different behavioral phenotypes based on avoidance behavior during re-exposure to a bobcat urine paired context (Albrechet-Souza & Gilpin, 2019; Albrechet-Souza et al., 2020; Edwards et al., 2013; Weera et al., 2020), anxiety and arousal behavior during an elevated-plus maze and acoustic startle response following TMT exposure (Schwendt et al., 2018), and anxiety-like behavior in an elevated plus maze and context avoidance behavior(Brodnik et al., 2017; Brodnik, Black, & Espana, 2020).

The size of the exposure chamber, addition of bedding to bottom of chamber, and inability to access the TMT odor, provides rats the opportunity to engage in a multitude of stress-coping behaviors. This is a unique characteristic of this specific model as it allows the engagement in different stress coping behaviors. Furthermore, by examining behavior during the stressor exposure, we can identify patterns of behaviors that may also be indicative of individual differences. For example, we can identify differences in the transition from defensive digging behavior to immobility and avoidance behavior or lack of transition between behaviors.

Overall, this predator odor exposure procedure detailed in this current protocol is a highly beneficial model to understand 1) behavioral changes indicative of a stress response, 2) quantify individual variability in behavioral response to stress and 3) how engagement in different behavioral coping strategies during TMT can predict long-term consequences of stress. This procedure is beneficial to understand neurobiological mechanisms associated with PTSD and comorbid disorders as well as producing behavioral profiles capable of modeling aspects of PTSD symptomology.

### Critical Parameters

#### Preventing contamination: Water and TMT chambers

It is critical that water and TMT exposure chambers are labeled as water or TMT to prevent contamination of water chamber with TMT. If at any time a water chamber is exposed to TMT (i.e. accidental pipette of TMT instead of water), that chamber should no longer be used as a water exposure chamber.

#### Reusing Chambers for Exposures (in same day)

During instances when there are not enough water and TMT chambers for the total number of rats needed for exposure, chambers may be reused. For example, consider an experiment that includes 24 animals, eight of which will be exposed to water and 16 that will be exposed to TMT. However, there are only six water chambers and 20 TMT chambers available. Chambers can be reused, but the bedding must be thrown out and chambers must be thoroughly cleaned (see Step 6. Cleaning under *Apparatus* section), before reusing. Chambers must also be dried before reassembling for next exposure. Proper cleaning procedures between exposures is necessary to prevent contamination of scents from animals, urine and fecal boli.

#### Bedding

All exposure chambers (water and TMT) should receive the same amount of bedding (600 ml). Differential amounts of bedding may affect the degree of engagement in the stress-reactive behaviors and affect classification of rats into TMT-subgroups. Furthermore, bedding used for exposure should be fresh, in the absence of any odors or previous contact with other rodents.

#### Exposure Environment

During exposure and recording, it is important to reduce noise and movements that could startle the animals or cause them to become anxious. Noises and movements can cause the animals to freeze, which would confound the quantification of freezing behavior produced by the exposure. To prevent noise and movements, it is recommended the researcher(s) leave the procedure room after exposure and recording has started. In addition, no one should enter the procedure room during exposure.

#### Recording

Camera set up is vital when recording exposures (*see Figure 3* for camera set up). For this protocol, exposures are recorded from the front view of the chambers. The camera should start recording prior to placing the first rat and chamber onto shelving unit. When place chambers onto shelving unit, the researcher should make every attempt to not block the recording view of rats that are already on the shelving unit. Also, cameras should be fully charged prior to starting the first recording. If the camera turns off during the recordings, that round of exposure videos will be affected and data will be lost..

### Troubleshooting

Common problems with the protocols, their causes, and potential solutions. Itemized in a 3- column table of Problem, Possible Cause, and Solution (see below for example).

**Table 1.** Troubleshooting.

| Problem | Possible Cause | Solution |
| --- | --- | --- |
| Control rats are exhibiting digging and immobility behavior that is similar to TMT rats and skewing data. | Control chambers have been contaminated by TMT. | Use new chambers for water exposures. If that is not an option, thoroughly clean chambers with Peroxiguard or 3% bleach solution. For best results, disassemble chambers (undo screws from walls) to scrub between walls and in corners. However, it is highly recommended that the control chambers that are contaminated stop being used for control group. |
| During TMT exposure, rats are obscured behind pile of bedding from digging, affecting scoring behavior (i.e. tracking and or view of rat). | This is normal digging behavior during TMT exposure. | If tracking for immobility and digging behavior is affected due to the pile of bedding rats formed during digging behavior, behavior should be scored by hand. If rats are digging behind bedding and are hidden, scorers should focus on the rodent's forepaws, shoulders and head looking for shoveling movements of bedding. This should be scored as digging behavior. |

#### Data Analysis

##### TMT Behavior

Analysis of total time spent engaging in stress-reactive behaviors (i.e. defensive digging, immobility behavior, avoidance, midline crossings, etc.) are first analyzed between water vs. TMT groups using individual t-tests. Analysis of behaviors across time (one minute segments) between water and TMT groups are analyzed using two way ANOVAs with time as a repeated factor. When separating into TMT-subgroups, groups are as follows: water, TMT-1 and TMT-2. Comparison of total time spent engaging in stress-reactive behaviors between groups is analyzed using a one-way ANOVA. Comparisons of stress-reactive behaviors across time between groups are analyzed by a two-way RM ANOVA with TMT exposure as a between-subjects factor and time as a within-subjects factor. Tukey multiple comparisons tests are used to follow up on significant main effects of groups and interactions.

##### Understanding Results

To provide representative results for this protocol, 24 female Long Evans rats (n=24) were used for water (n=8) or TMT (n=16) exposure and the procedure described above.

###### Water vs. TMT Exposure Data

We first analyzed digging and immobility behavior (total and across time) between water and TMT exposed rats.

##### Digging Behavior

During TMT exposure, rats engaged in significantly more digging behavior compared to controls (Figure 4A; unpaired t-test; *p* < 0.05). In addition, a two-way ANOVA for repeated measures (RM) showed a significant digging x time interaction (Figure 4B,*F* (9,216) = 6.12, *p* < 0.05) such that rats exposed to TMT engaged in greater digging behavior during minutes 2-8 of exposure compared to controls.

**Figure 4.**
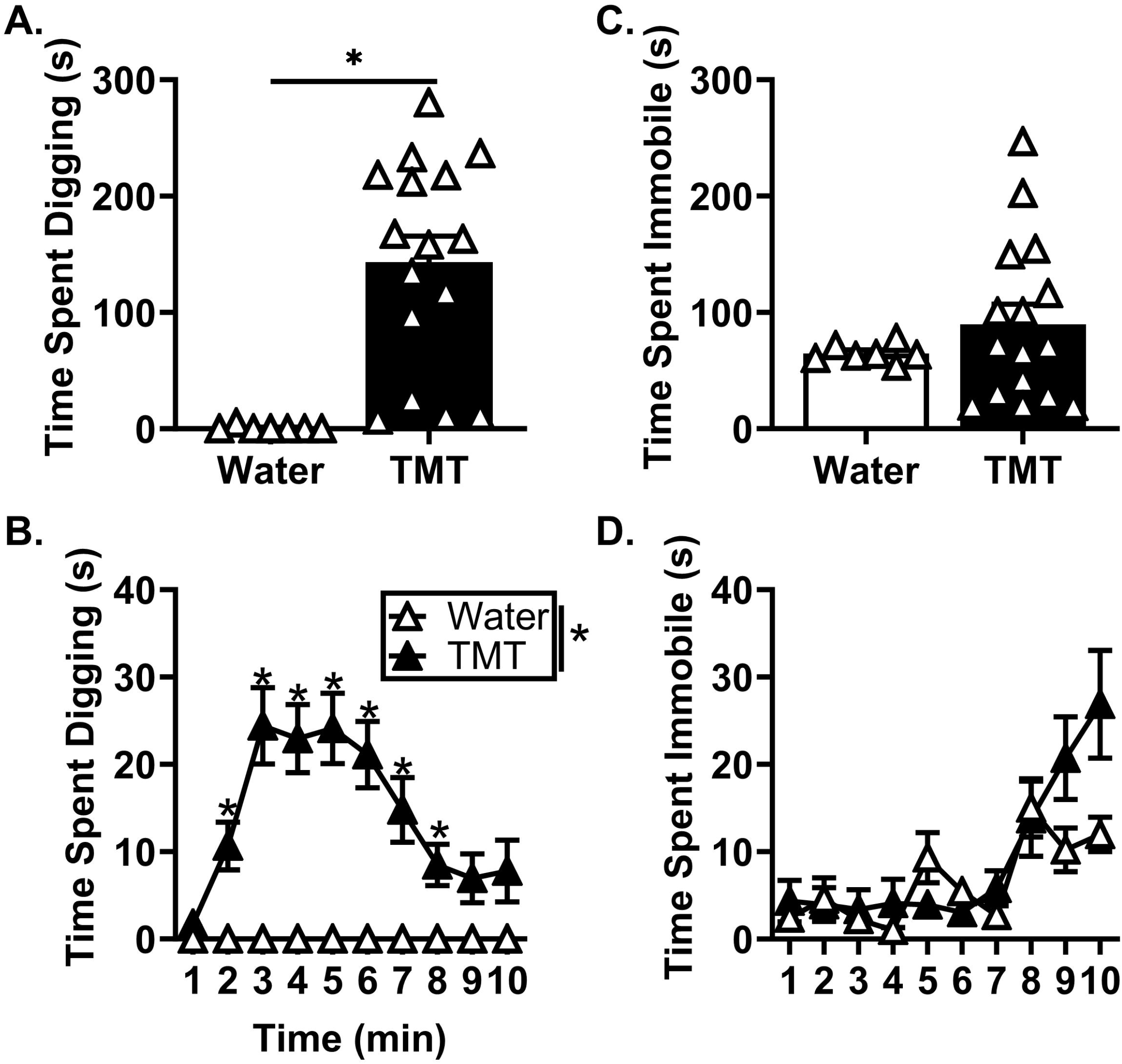
Effects of TMT exposure on stress-reactive behaviors. Female rats exposed to TMT spent significantly greater total time (**A**) and across time (**C**) engaging in digging behavior compared to controls. Female rats exposed to TMT did not show differences in total time spent immobile (**B**) or across time (**D**) compared to controls. **\****p* < 0.05 vs. controls. Mean ± SEM

##### Immobility Behavior

During TMT exposure, rats exposed to TMT did not show a difference in total time spent immobile compared to controls (Figure 4C; unpaired t-test;*p* > 0.05). In addition, there were no significant immobility x time interaction or main effects of TMT exposure on immobility behavior (Figure 4D, *p* > 0.05). However, the spread in total time immobile in TMT exposed rats varied between 20s to 240s, suggesting a large range in variability in low and high immobile rats. This is an important observation as it is pertains to individual differences, that will be visible when analyzing these data across TMT-subgroups.

###### Water vs. TMT-1 vs. TMT-2 Subgroups Exposure Data

Next, we analyzed digging and immobility behavior (total and across time) between controls and TMT-subgroups. Rats were classified into TMT subgroups by a digging/immobility ratio score based on their behavior during the TMT exposure (Figure 5A,*see representative schematic*). Representation of digging/immobility (D/I) ratios in female TMT-subgroups 1 and 2 is illustrated in Figure 5B, showing rats with a D/I ratio of less than 1.0 as TMT-1 and rats with a D/I ratio greater than 1.0 as TMT-2.

**Figure 5.**
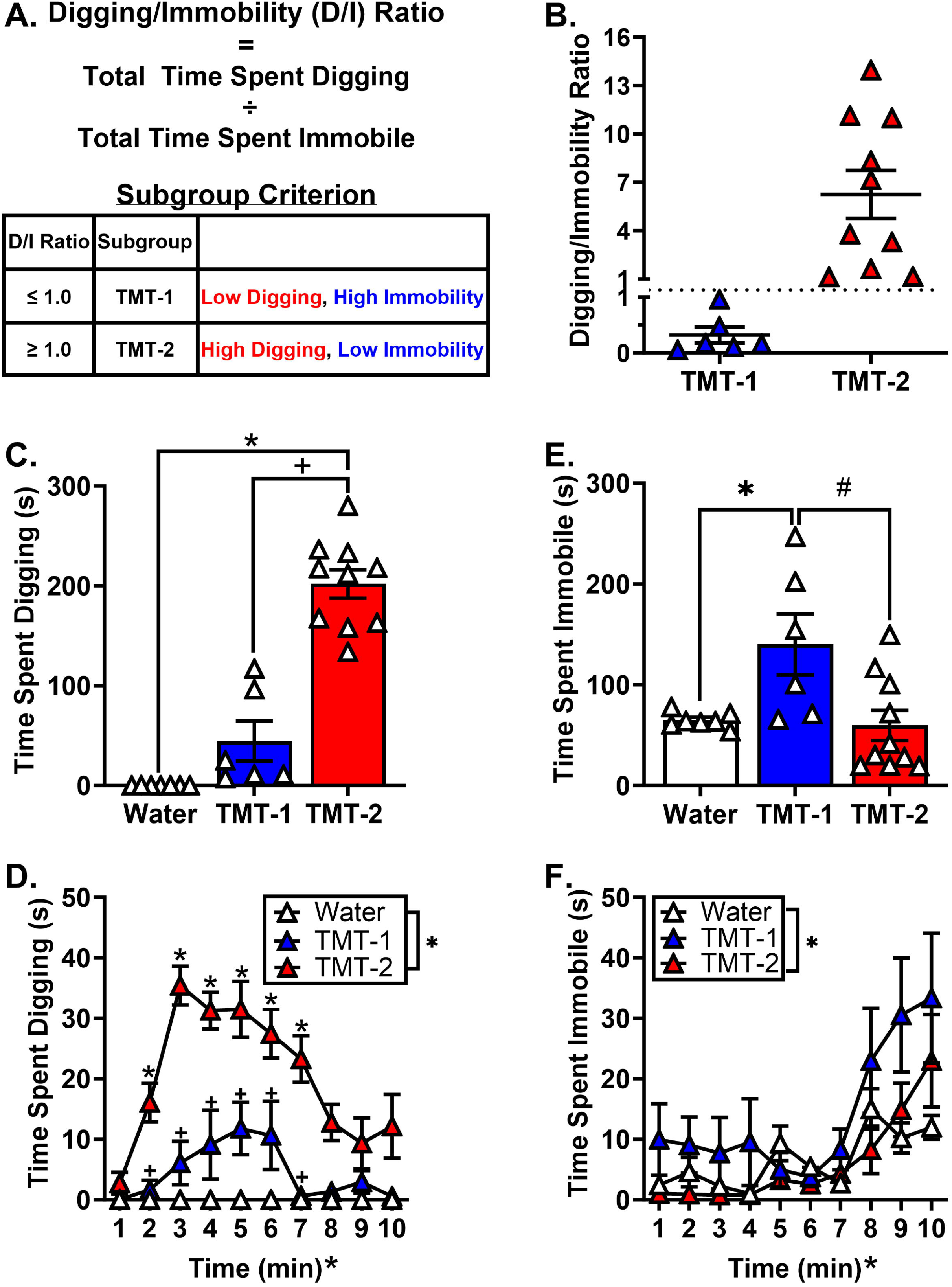
Formation of TMT-subgroups and behavioral phenotype data. (**A**) Description of digging/immobility ratio score and subgroup criterion. (**B**) Representation of digging/immobility scores in female TMT-groups 1 and 2. Female rats in TMT-2 engaged in significantly greater digging behavior compared to controls and TMT-1 (**C**), as well as across 10 min TMT exposure (**D**). Female rats in TMT-1 engaged in significantly greater immobility behavior compared to controls and TMT-2 (**E**) but not across time (**F**). **\****p* < 0.05 vs. controls; **#** *p* < 0.05 vs. TMT-2, **+** *p* < 0.05 vs. TMT-1. Mean ± SEM

##### Digging Behavior

One-way ANOVA showed a significant main effect of exposure (Figure 5C,*F* (2,20) = 63.76, *p* < 0.05), such that TMT-2 rats engaged in greater total time spent digging compared to water and TMT-1 subgroups. In addition, when analyzing digging behavior across time, a two-way RM ANOVA showed a significant exposure x time interaction (Figure 5D,*F* (18,180) = 4.08, *p* < 0.05), such that rats in TMT-2 subgroup engaged in greater digging behavior compared to controls and TMT-1 subgroup from minute 2 through 7 (*p* ≤ 0.05). There was also a significant main effect of exposure (Figure 5D, *F* (2,20) = 63.76, *p* < 0.05) and time (Figure 5D, *F* (9,180) = 7.61, *p* < 0.05).

##### Immobility Behavior

An one-way ANOVA showed a significant main effect of exposure (Figure 5E, *F* (2,20) = 5.70, *p* < 0.05, such that TMT-1 subgroup engaged in greater immobility behavior compared to controls and TMT-subgroup 2. When analyzing immobility behavior across time, there was no significant interaction between exposure x time (Figure 5F,*p* > 0.05); therefore, we cannot compare immobility behavior between exposure groups at individual minute time points. There was a main effect of exposure (*F* (2,20) = 5.70, *p* < 0.05) and time (*F* (9,180) = 10.43, *p* < 0.05). Therefore, when separating animals into subgroups, we show that rats in TMT- 1 engage in high immobility behavior during TMT exposure.

##### Time Considerations

Time considerations for predator odor stressor exposure using TMT (Basic Protocol 1) is split into three parts: 1) preparation, 2) exposure and 3) cleaning.

1. Preparation time to set up chambers for exposure will depend on how many animals are being exposed. There must be ample time for organizing the appropriate number of water and TMT chambers needed for exposure, adding bedding to the chambers, placing metal basket and holder to chambers, adding filter paper to the baskets and securing lids on chambers.
2. Experimenters should prepare for at least 30 minutes for each individual exposure due to the amount of time it takes to: organize chambers under fume hood to prep for water or TMT pipetting, transport rats to procedure room, pipette water or TMT onto each filter paper, place rats into assigned chamber and onto shelving unit for recording, and lastly, the 10-minute exposure time. Multiple groups of rats can be run within a single day.
3. Cleaning the chambers and procedure room after TMT exposure can take a considerable amount of time. Depending how many chambers were used for exposures, cleaning can take up to 1-2 hours and duration of all exposures for the day. Chambers should be cleaned first to allow for deep cleaning and air drying on wire rack. For procedure room, wiping down all surfaces, sweeping, mopping and throwing out trash should be completed second.

## Supporting information

Biorender publication licenses for figures

## ACKNOWLEDGMENTS

This work was supported in part by the National Institute of Health AA029730 (LCO) and AA026537 (JB) and by the Bowles Center for Alcohol Studies. Illustrations of water/TMT exposures and exposure chambers created with BioRender.com.

## AUTHOR CONTRIBUTIONS

**Laura C. Ornelas:** Conceptualization, Data curation, Formal analysis, Investigation, Methodology, Validation, Visualization, Roles/Writing – original draft, Writing – review & editing. **Joyce Besheer:** Conceptualization, Funding acquisition, Investigation,

Methodology, Project administration, Supervision, Validation, Visualization, Roles/Writing - original draft, Writing - review & editing.

## STATEMENT ON THE WELFARE OF ANIMALS

Animals were under continuous care and monitoring by veterinary staff from the Division of Comparative Medicine at UNC-Chapel Hill. All procedures were conducted in accordance with the NIH Guide for the Care and Use of Laboratory Animals and institutional guidelines.

## DATA AVAILABILITY STATEMENT

Data available on request from the authors.

## LITERATURE CITED

Alavi, M., Ryabinin, A. E., Helms, M. L., Nipper, M. A., Devaud, L. L., & Finn, D. A. (2022). Sensitivity and Resilience to Predator Stress-Enhanced Ethanol Drinking Is Associated With Sex-Dependent Differences in Stress-Regulating Systems.Front Behav Neurosci, 16, 834880. doi:10.3389/fnbeh.2022.834880

Albrechet-Souza, L., & Gilpin, N. W. (2019). The predator odor avoidance model of post-traumatic stress disorder in rats. Behav Pharmacol, *30*(2 and 3-Spec Issue), 105-114. doi:10.1097/FBP.0000000000000460

Albrechet-Souza, L., Schratz, C. L., & Gilpin, N. W. (2020). Sex differences in traumatic stress reactivity in rats with and without a history of alcohol drinking.Biol Sex Differ, 11(1), 27. doi:10.1186/s13293-020-00303-w

APA. (2013). Diagnostic and statistical manual of mental disorders (5th ed.). Washington, DC.

Becker, H. C., Lopez, M. F., King, C. E., & Griffin, W. C. (2023). Oxytocin Reduces Sensitized Stress-Induced Alcohol Relapse in a Model of Posttraumatic Stress Disorder and Alcohol Use Disorder Comorbidity. Biol Psychiatry, 94(3), 215–225. doi:10.1016/j.biopsych.2022.12.003

Brodnik, Z. D., Black, E. M., Clark, M. J., Kornsey, K. N., Snyder, N. W., & Espana, R. A. (2017). Susceptibility to traumatic stress sensitizes the dopaminergic response to cocaine and increases motivation for cocaine. Neuropharmacology, 125, 295–307. doi:10.1016/j.neuropharm.2017.07.032

Brodnik, Z. D., Black, E. M., & Espana, R. A. (2020). Accelerated development of cocaine-associated dopamine transients and cocaine use vulnerability following traumatic stress. Neuropsychopharmacology, 45(3), 472–481. doi:10.1038/s41386-019-0526-1

Center for Substance Abuse, T. (2014). SAMHSA/CSAT Treatment Improvement Protocols Trauma-Informed Care in Behavioral Health Services. Rockville (MD): Substance Abuse and Mental Health Services Administration (US).

Cohen, H., Kozlovsky, N., Alona, C., Matar, M. A., & Joseph, Z. (2012). Animal model for PTSD: from clinical concept to translational research. Neuropharmacology, 62(2), 715–724.

De Boer, S. F., & Koolhaas, J. M. (2003). Defensive burying in rodents: ethology, neurobiology and psychopharmacology. Eur J Pharmacol, 463(1-3), 145–161. doi:10.1016/s0014-2999(03)01278-0

Edwards, S., Baynes, B. B., Carmichael, C. Y., Zamora-Martinez, E. R., Barrus, M., Koob, G. F., & Gilpin, N. W. (2013). Traumatic stress reactivity promotes excessive alcohol drinking and alters the balance of prefrontal cortex-amygdala activity.Transl Psychiatry, 3, e296. doi:10.1038/tp.2013.70

King, C. E., & Becker, H. C. (2019). Oxytocin attenuates stress-induced reinstatement of alcohol seeking behavior in male and female mice. Psychopharmacology (Berl), 236(9), 2613–2622. doi:10.1007/s00213-019-05233-z

Makhijani, V. H., Franklin, J. P., Van Voorhies, K., Fortino, B., & Besheer, J. (2021). The synthetically produced predator odor 2,5-dihydro-2,4,5-trimethylthiazoline increases alcohol self-administration and alters basolateral amygdala response to alcohol in rats. Psychopharmacology (Berl), 238(1), 67–82. doi:10.1007/s00213-020-05659-w

Ornelas, L. C., Tyler, R. E., Irukulapati, P., Paladugu, S., & Besheer, J. (2021). Increased alcohol self-administration following exposure to the predator odor TMT in active coping female rats. Behav Brain Res, 402, 113068. doi:10.1016/j.bbr.2020.113068

Ornelas, L. C., Van Voorhies, K., & Besheer, J. (2021). The role of the nucleus reuniens in regulating contextual conditioning with the predator odor TMT in female rats. Psychopharmacology (Berl). doi:10.1007/s00213-021-05957-x

Rosen, J. B., Donley, M. P., Gray, D., West, E. A., Morgan, M. A., & Schulkin, J. (2008). Chronic corticosterone administration does not potentiate unconditioned freezing to the predator odor, trimethylthiazoline. Behav Brain Res, 194(1), 32–38. doi:10.1016/j.bbr.2008.06.019

Rosen, J. B., West, E. A., & Donley, M. P. (2006). Not all rat strains are equal: differential unconditioned fear responses to the synthetic fox odor 2,4,5-trimethylthiazoline in three outbred rat strains. Behav Neurosci, 120(2), 290–297. doi:10.1037/0735-7044.120.2.290

Schwendt, M., Shallcross, J., Hadad, N. A., Namba, M. D., Hiller, H., Wu, L., Krause, E. G., & Knackstedt, L. A. (2018). A novel rat model of comorbid PTSD and addiction reveals intersections between stress susceptibility and enhanced cocaine seeking with a role for mGlu5 receptors. Transl Psychiatry, 8(1), 209. doi:10.1038/s41398-018-0265-9

Staples, L. G. (2010). Predator odor avoidance as a rodent model of anxiety: learning-mediated consequences beyond the initial exposure.Neurobiology of learning and memory, 94(4), 435–445.

Tyler, R. E., Bluitt, M. N., Engers, J. L., Lindsley, C. W., & Besheer, J. (2022). The effects of predator odor (TMT) exposure and mGlu(3) NAM pretreatment on behavioral and NMDA receptor adaptations in the brain. Neuropharmacology, 207, 108943. doi:10.1016/j.neuropharm.2022.108943

Tyler, R. E., Weinberg, B. Z. S., Lovelock, D. F., Ornelas, L. C., & Besheer, J. (2020). Exposure to the predator odor TMT induces early and late differential gene expression related to stress and excitatory synaptic function throughout the brain in male rats.Genes Brain Behav, 19(8), e12684. doi:10.1111/gbb.12684

Verbitsky, A., Dopfel, D., & Zhang, N. (2020). Rodent models of post-traumatic stress disorder: behavioral assessment. Transl Psychiatry, 10(1), 132. doi:10.1038/s41398-020-0806-x

Weera, M. M., Schreiber, A. L., Avegno, E. M., & Gilpin, N. W. (2020). The role of central amygdala corticotropin-releasing factor in predator odor stress-induced avoidance behavior and escalated alcohol drinking in rats. Neuropharmacology, 166, 107979. doi:10.1016/j.neuropharm.2020.107979

